# Uneven Walking is Challenging: step-ups or extended steps that is the question

**DOI:** 10.1101/2024.07.10.602932

**Authors:** Seyed-Saleh Hosseini-Yazdi

**Author notes:** **Correspondence Address:** Seyed Saleh Hosseini Yazdi, Biomedical Engineering Department, Schulich School of Engineering, University of Calgary, 2500 University Dr NW, Calgary AB T2N1N4.

## Abstract

Uneven terrain presents significant challenges for walkers, resulting in increased energy expenditures. Given that Center of Mass (COM) work reflects this energy demand, it's reasonable to assume that individuals also seek strategies to minimize mechanical work. One such strategy involves deciding between extending step length to avoid terrain irregularities or simply traversing over them. Each approach carries its own mechanical cost, leading to the adoption of the less costly option. To investigate this, we conducted a simulation focusing on COM mechanical work under the assumption that gait energy is entirely provided through pre-emptive push-off. We examined the COM work required for step length extension, ranging from nominal to twice its magnitude, and compared it with the mechanical work needed for step-ups from zero to 0.05 m. The simulation revealed a critical threshold for a given walking velocity and perturbation amplitude: below it, extending step length was more favorable, while beyond it, landing atop perturbations became the preferred choice. As perturbation amplitude rose, the magnitude of the threshold also increased.

## Introduction

Walking energetics is one of the main determinants of gait. It is suggested that humans nominal gait coincides with the minimum metabolic cost [1]. However, when faced with challenging walking conditions, various factors also come into play, influencing our gait [2]. One challenge of navigating uneven terrain lies in the increased effort required to overcome gravity during step-ups, as well as the heightened collision dissipations during step-downs, both of which amplify the mechanical work demands [3], impacting the overall energetics of walking [4]. Consequently, the parameters of our gait on uneven surfaces likely reflect our efforts to minimize mechanical work. During each step-to-step transition, humans evaluate different options to choose the most metabolically efficient. Whether it's navigating over a terrain perturbation or adjusting the step length to avoid it, perhaps humans instinctively opt for the least mechanically costly option. The mechanical cost possibly is the deriver to decide on the favorable option.

In complex landscapes like uneven terrains, step variabilities tend to rise [2]. This increase in step variability is believed to be a result of frequent corrections made to sustain the gait amid challenging walking conditions [1]. Consequently, heightened step length variabilities may signify a trade-off between one foot landing location versus another, beyond just maintaining the balance, and maybe minimizing mechanical work during walking.

During walking, the process of transitioning from one stance leg to another, as highlighted by Kuo et al. [5], involves simultaneous positive and negative works [6]. These works serve to align the trajectory of the Center of Mass (COM) with the next stance [7]. According to Donelan et al. [4], the positive work during this transition roughly mirrors the energetic demands of walking. Additionally, Kuo [8] proposed that the work during the step transition is influenced by the average walking velocity and the angle between the leading and trailing legs at the transition point. It's been suggested that the positive active work results from a preemptive push-off impulse (*po* = *ν*_*ave*_ ⋅ tan *α*) at toe-off, which increases with the fourth power of the step length beyond its nominal value [4,8]. Hence, it is plausible to consider the push-off work as a key determinant for establishing the upper limit of step length.

The lower bound of the human step length is also restricted by the increasing cost associated with limb swing or step frequency [9,10]. It is suggested that for a given walking velocity, there exists a preferred step length and step frequency [11,12]. However, if the step frequency exceeds its nominal value, the force rate generation cost of limb swing begins to dominate the walking energetics [10], and this cost escalates substantially with the fourth power of the step frequency [9]. This phenomenon effectively limits humans from taking short steps [5]. Consequently, during uneven walking, shorter step lengths may be viewed as transitory adjustments made to regulate COM movements.

Walking against gravity also poses a significant energetic challenge[3], leading humans to potentially avoid or minimize step elevation changes whenever feasible. One possible strategy is to vary step lengths to navigate around or over the terrain obstacles. However, the increased mechanical cost associated with longer steps may prompt individuals to choose another approach, such as traversing atop terrain complexities (Figure 1). In essence, we propose the development of a mechanistic metric to determine when humans might opt for longer step lengths versus landing atop the terrain perturbations, considering the associated mechanical work against gravity. The decision-making process could hinge on comparing the magnitudes of mechanical costs to select the most energy-efficient strategy.

**Figure 1.**
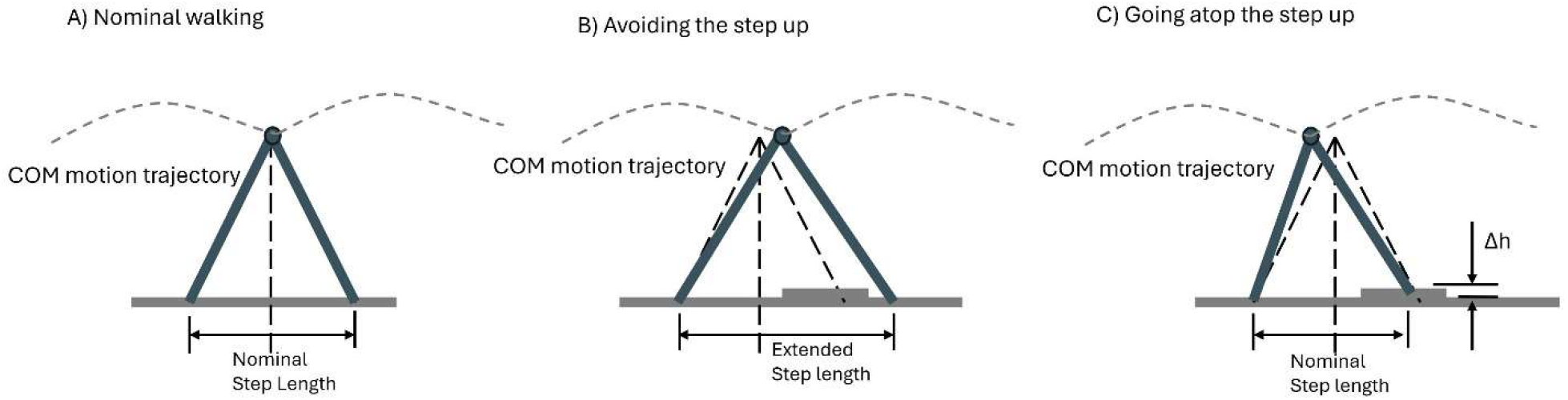
Schematics representing (A) nominal gait for a given walking speed over the even terrain, (B) the perturbation avoidance by extending the step length, (C) landing atop the perturbation when perturbation avoidance is not possible.

## Materials and Methods

We utilized the powered simplest walking model method [8,13] to estimate the mechanical cost of the step-to-step transition based on the step length and work against gravity to sustain the average walking speed. For a step up, the optimum pre-emptive push-off provides the required mechanical energy to sustain the average walking speed (*ν*_*ave*_) and work against the gravity (*g* ⋅ Δ*h*). As such, for any step-to-step transition, the post transition walking speed 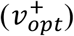 magnitude must present the required kinetic energy. We considering the pre-transition velocity as *ν*^−^ = ν_*ave*_ [8], the mid transition velocity magnitude and its angle with the pre-transition velocity for a given push-off can be calculated as (Figure 2):

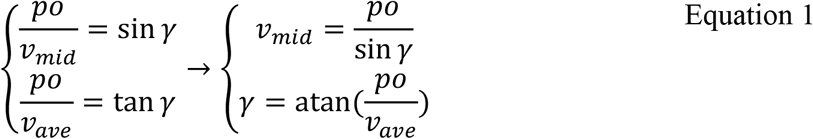

The resulted post transition velocity becomes:

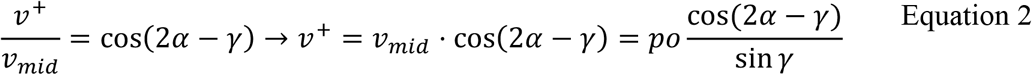

We simplify 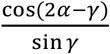as 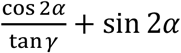, therefore the post transition velocity becomes:

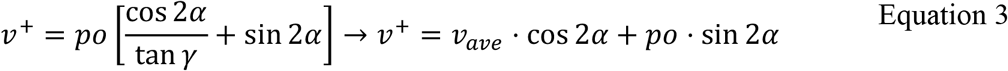

The post transition velocity must provide the average kinetic energy 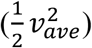 and cover the work against the gravity (*g* ⋅ Δ *h*):

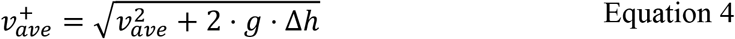

Therefore, the optimal push-off impulse becomes:

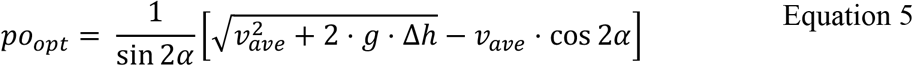

**Figure 2.**
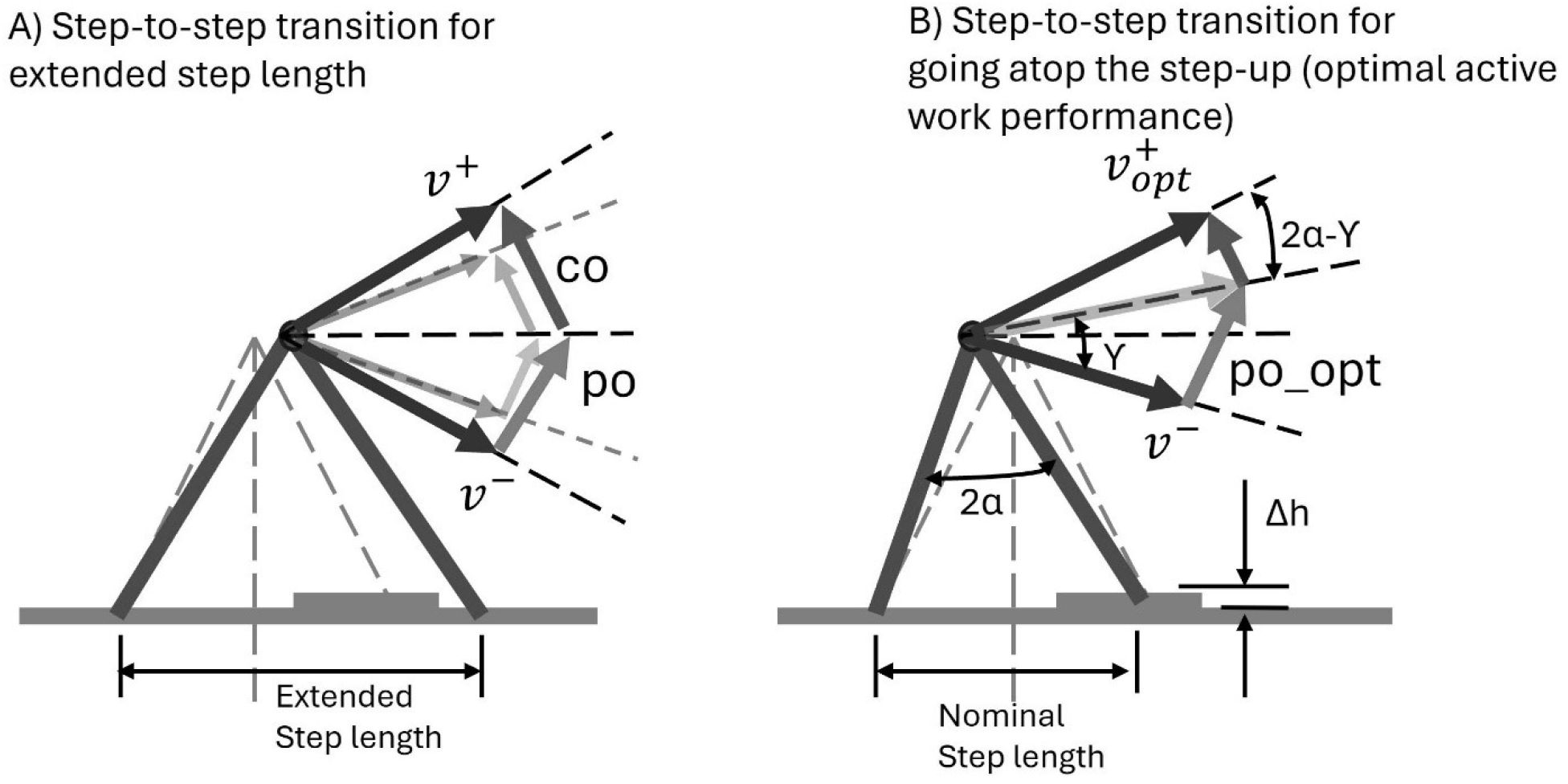
Step to step transition (A) when the step length is extended. As such, the push-off and collision works grow. The Solid line push-off (po) and collision (co) are for longer steps whereas the shaded impulses are for the nominal walking. (B) The optimal pre-emptive push-off and post transition velocity to sustain the average walking speed and compensate for the gravity work.

## Results

The angle between the leading and trailing leg at the transition point (2α) represented the step length under the assumption of unity leg length. Consequently, an increase in this angle corresponded to an increase in step length. Assuming a walking speed of 1.25 m ⋅ s^−1^, we assumed α to be 0.35 radians [14]. We increased α from 0.35 to 0.70 radians that resulted in a step length increase from 0.700 m to 1.343 m, or +91.9%, the associated pre-emptive push-off ranged from 0.104 J ⋅ kg^−1^ to 0.554 J ⋅ kg^−1^ (+432.7%, Figure 3A & B).

**Figure 3.**
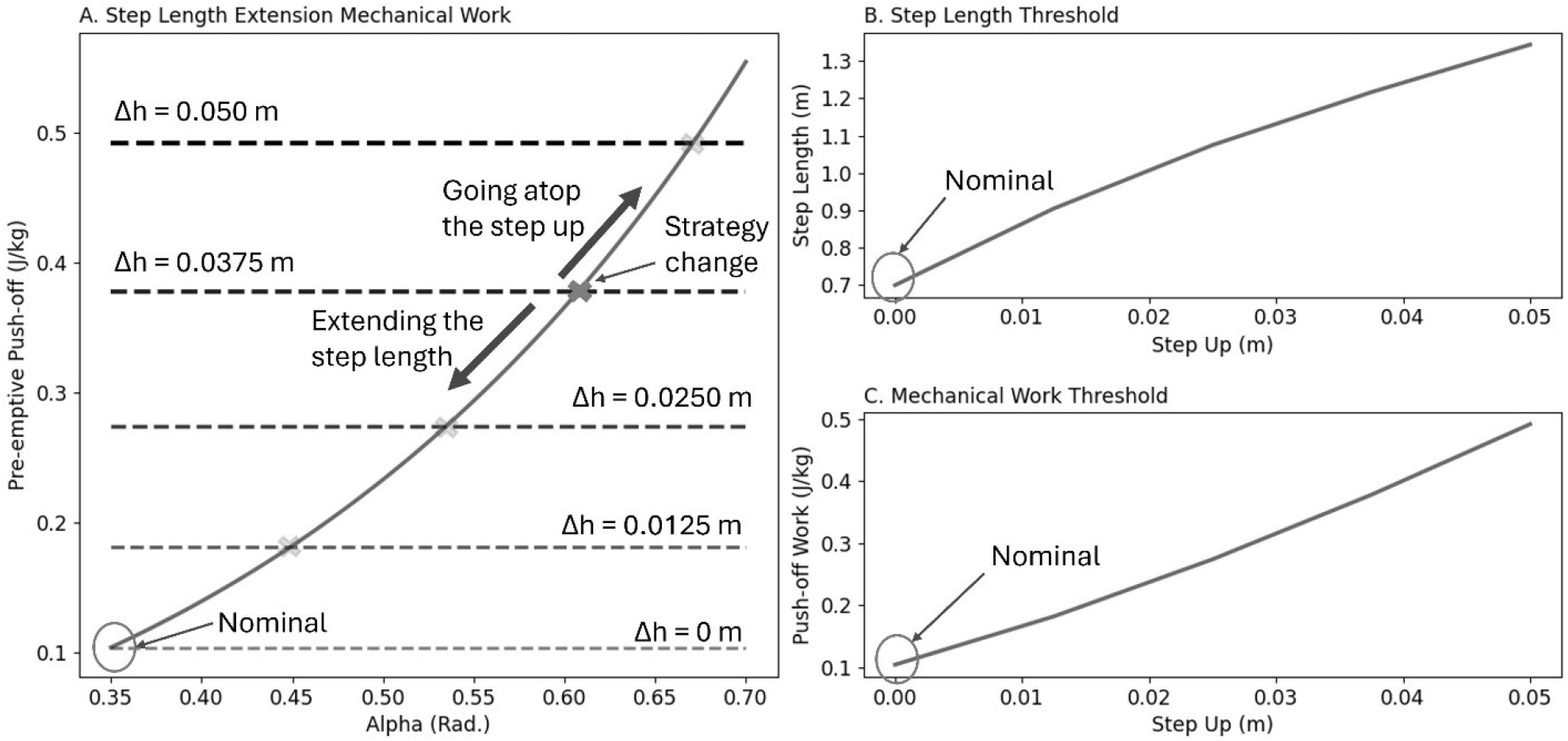
(A) analytical COM work for the extended steps (v = 1.25 *m* ⋅ *s*^−1^). The transition work increases with the step length rise. Humans also incur large mechanical cost to perform works against gravity. (B) Step length increase threshold for step-ups. With the rise of the step-up amplitudes, it becomes mechanically viable to extend the step length more. (C) The mechanical work threshold for step-ups. With the increase of step-up amplitude, the mechanical work threshold to switch from step length extension to going atop the perturbation also rises.

It was suggested that walking mechanical economy was maximized when all the energy required for the gait was provided entirely by the pre-emptive push-off [5]. Therefore, we also assumed that the optimal pre-emptive push-off covered the energy required to work against gravity. During step-ups, where step lengths were maintained at nominal values, when the step-up elevation increased from zero (even) to 0.050 m, the optimal pre-emptive push-offs ranged from 0.104 J ⋅ kg^−1^ to 0.492 J ⋅ kg^−1^ (+373.1%, and Figure 3C).

## Discussion

It has been observed that during uneven walking, humans deviate from their nominal gait [2], adapting to the perturbations of complex terrains. Alongside other goals like maintaining balance, it's important for humans to also manage their walking energetics efficiently [15]. To adjust to terrain complexities, humans make trade-offs, incurring some metabolic costs to avoid larger energy expenditures [16]. The variability in step length during uneven walking, whether taking longer or shorter steps, may reflect these decisions. Calculating the COM step-to-step transition work and the work against gravity could offer a mechanistic metric for determining the upper limit of step length.

Our simulation has depicted that at a given walking velocity, there exists a threshold below which it is more economical to take longer steps. This threshold signifies the point where the mechanical cost of stepping up and extending step length become equal. Beyond this threshold, it must be more favorable for humans to take a nominal step and navigate atop the perturbation, as the cost of performing gravity work is less than that of extending step length (Figure 4).

**Figure 4.**
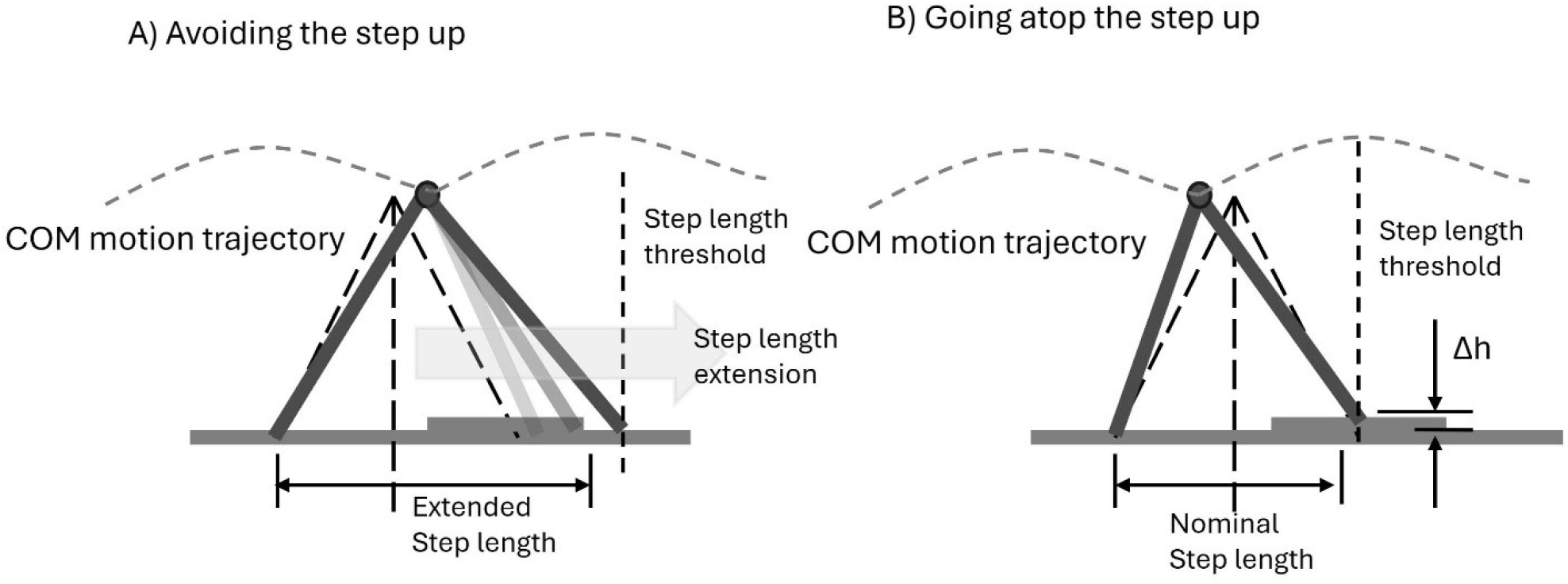
(A) With the length of the terrain perturbation (step up), humans may extend their legs to avoid the associated gravity work by avoiding the perturbation. (B) There is an upper limit for the step length extension beyond which the mechanical cost of going atop the perturbation becomes less than the perturbation avoidance.

The model presented here does not provide an estimation for a lower bound on step length. It's conceivable that scenarios exist where taking shorter steps and compensating for the lost time in subsequent steps could be energetically more efficient. However, determining the lower step length is most likely influenced by the muscle force generation cost, which is primarily governed by step frequency [10]. Therefore, a separate model would be required to address this aspect.

The presented model has some other limitations that need to be addressed for more accurate results. Firstly, the assumption of universal leg length (1.0 m) does not reflect the diversity of human leg lengths, which vary among individuals. Additionally, the range of motion at the hip joint imposes a limit on achievable step lengths, making excessive step lengths unattainable. Furthermore, the simulation assumes a return to even terrain beyond perturbation. However, in reality, subsequent steps may encounter different elevations, altering the optimal step length threshold (Figure 5). A step up following a perturbation may negate the advantage of perturbation avoidance, while a step down could increase flight time duration that increases the exerted push-off work magnitude [5]. In turn, it increases the metabolic cost of walking [4]. Therefore, the perturbation avoidance advantage possibly diminishes. As vision contributes to the selection of the subsequent foot landing [17], the lookahead horizon may also play a crucial part in assessment of the uneven terrain negotiation [18]. Finally, the simulation doesn't penalize lifting limbs to avoid perturbations. Humans may increase feet clearance to clear obstacles, resulting in greater work against gravity (peripheral work). This additional effort energetics during perturbation avoidance may further limit the upper limit of step length extension. Addressing these limitations would enhance the model's accuracy in simulating human walking behavior on uneven terrain.

**Figure 5.**
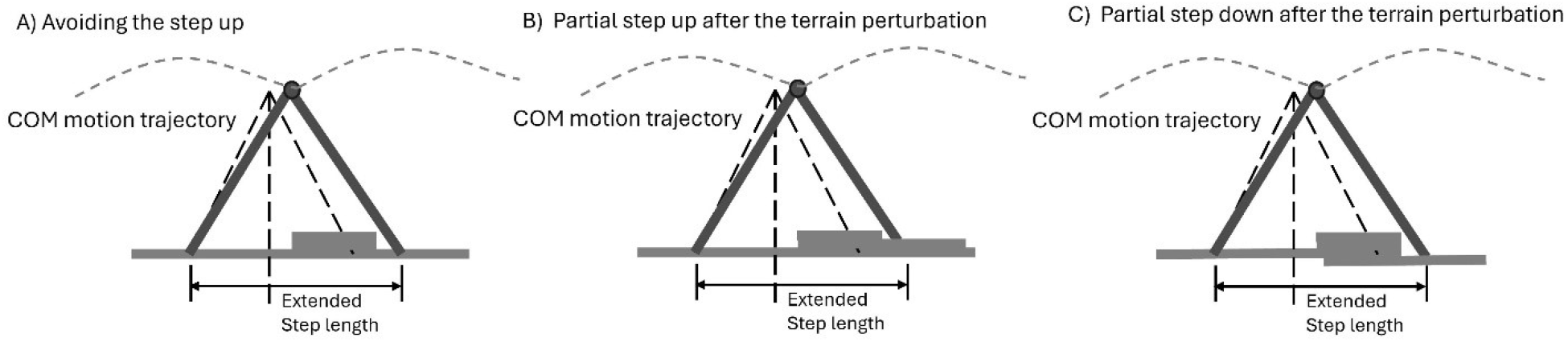
(A) Landing on the same elevation after the terrain perturbation avoidance, (B) partial step up after the terrain perturbation, (C) partial step down after the terrain perturbation.

In summary, humans exhibit complex coordination of various gait parameters during walking [19] to achieve balance [1], accomplish tasks [20], and conserve metabolic energy [21]. Uneven terrain introduces higher step variabilities [2], suggesting trade-offs aimed at also minimizing energy expenditure through perturbation avoidance or negotiation. Our simulation reveals a critical threshold for COM work, below which extending step length is more cost-effective, while above it, navigating atop perturbations becomes preferable. However, our model has limitations. It doesn't fully account for individual leg lengths or joint ranges of motion, which can constrain step length. Additionally, perturbation avoidance may incur metabolic costs from lifting limbs to clear obstacles. To understand these factors better, experimental studies with varied perturbation amplitudes and lengths are necessary to determine how humans adapt their strategies. Such research can provide valuable insights into the interplay between gait dynamics and terrain negotiation in human locomotion.

